# PTPRC, KDM5C, GABBR1 and HDAC1 are the major targets of valproic acid in regulation of its anticonvulsant pharmacological effects

**DOI:** 10.1101/2022.12.12.520029

**Authors:** Arun HS Kumar

## Abstract

**Background:** Valproic acid (VPA) is a small molecule which is the 3^rd^ most prescribed drug among anticonvulsant therapeutics. Understanding of the pharmacology of VPA targets will help optimally rationalise the therapeutic effects and also minimise the undesired outcomes. Hence this study analysed the human specific targets of VPA and assessed the affinity of VPA to these targets to interpret potential safe therapeutic range for VPA.

**Materials and Methods:** The targets of VPA were identified from the SwissTargetPrediction server and STITCH database and analysed for their affinity with VPA using Autodock vina 1.2.0. The volume of distribution (Vd, L) and the dose of VPA reported in the DrugBank database was used for estimation of the plasma and CSF concentration. The plasma and CSF Concentration Affinity (CA) ratio of VPA against each of the high affinity targets was assessed at variable Vd (0.1 to 0.4 L/kg) to identify the therapeutic safety window of VPA.

**Results:** The plasma/CSF concentration of VPA range from 170 to 7000 µM and 17 to 700 µM respectively. The plasma concentration achieved was within the safety limits (170 to 700 µM) at higher Vd (>10 L), while at lower Vd (<10L), the plasma or CSF concentration achieved was of concern at VPA dose of >1000 mg/day. The plasma concentration at very low Vd (< 2L) was of concern even at dose of 500 mg/day. The affinity of VPA against all its human specific targets ranged from 2.9 to 52.1 mM. The CA ratio of VPA against its high affinity target was observed to be greater than 0.8, indicating potentially significant modulation of these targets. The following four targets showed CA ratio of over 1: PTPRC, KDM5C, GABBR1 and HDAC1, indicating their preferential targeting by VPA. CES1 and SLC22A12 are high affinity targets of VPA which can contribute to its undesired pharmacological effects (CNS oedema and hepatotoxicity).

**Conclusion:** This study offers a novel insight into the anticonvulsant and undesired pharmacology of VPA by specifically identifying the targets involved and recommends an evidence based approach to personalise dose titration of VPA to achieve optimal therapeutic benefits.

## Introduction

Valproic acid (VPA) is a small molecule which is approved or used off label for the treatment of epilepsy, migraine, bipolar disorder/mania and depression.^[1-4]^ Ironically the precise mechanism/s by which the therapeutic benefits of VPA are achieved is not known. Some of the major pharmacological effects of VPA are by 1) increasing the GABA levels in CNS (amygdala and hippocampus), 2) modulating ion channels (sodium, potassium and/or calcium) and 3) inhibition of histone deacetylase’s.^[3, 5]^ The effects of VPA in increasing GABA levels is shown to be achieved through inhibition of GABA transaminase and succinate semialdehyde dehydrogenase, as both these enzymes are involved in degradation of GABA to succinate.^[2, 3, 5]^ With over 6 million prescriptions per year in USA, VPA ranks 109^th^ among top drug prescriptions and is 3^rd^ most prescribed drug among anticonvulsant therapeutics (https://clincalc.com/). Despite its extensive use in clinical practice, the lack of clarity on its therapeutic target/s is of concern. While several targets have been reported for VPA based on studies using animal models, the relevance of these targets to the pharmacodynamic effects of VPA in humans is unclear.^[5, 6]^ While many of these targets support the favourable therapeutic effects of VPA in treating epilepsy, migraine, bipolar disorder/mania and depression^[3-5]^ an equal number of other targets highlight the undesirable outcomes of VPA use.^[2, 7]^ While such counter pharmacodynamic features are not unique to VPA and paraphs most drugs have such diverse effects,^[8, 9]^ the important pharmacological factor which influences desired pharmacodynamic effect is the dose of the drug used in relation of the affinity of the drug to its target. Evaluation of the affinity of VPA to all its targets helps optimally rationalise the pharmacological outcomes from therapeutic use of VPA. Hence this study analysed the human specific targets of VPA and assessed the affinity of VPA to these targets to interpret potential safe therapeutic range for VPA.

## Materials and Methods

The targets of valproic acid (VPA) were identified from the SwissTargetPrediction server and STITCH database as reported before ^[8-11]^ and analysed for their affinity with VPA using Autodock vina 1.2.0. Briefly, the isomeric SMILES sequence of the VPA obtained from the PubChem database was inputted into the SwissTargetPrediction server and STITCH database to identify the targets specific to homo sapiens.^[8, 9, 11]^ The PDB structures of all the target proteins (n=110) and VPA were processed for docking in Autodock vina 1.2.0 for the estimation of affinity value. The volume of distribution (L) and the dose of VPA reported in the DrugBank database and NHS drug formulary^[1]^ respectively was used for estimation of the plasma and CSF concentration (µM) of VPA achievable.^[12, 13]^ The concentration of VPA in the CSF was calculated on the basis of 10% of plasma concentration as widely reported in the literature.^[14, 15]^

The affinity of VPA to all its targets was arranged in ascending order and all high affinity targets (Affinity < 5 mM) were selected for the analysis of concentration affinity ratio (CA ratio).^[8]^ 2-en-VPA and 3-keto-VPA are reported to be the most prevalent metabolites^[3, 5]^ of VPA and the affinity of these two metabolites against all high affinity targets was also assessed and compared to that of VPA. The plasma and CSF CA ratio of VPA against each of the high affinity targets was further assessed at variable volume of distribution (0.1 to 0.4 L/kg) values. The CA ratio is presented as mean ± SD of the values estimated from low to high range of volume of distribution.

The protein expression profile of all high affinity targets of VPA were text mined from the Human Protein Atlas database (https://www.proteinatlas.org) and compared.^[9, 11]^ For the analysis in this study only high protein expression values was considered while the medium and low expression tissues/organs were ignored.

## Results

Valproic acid (VPA) is used in the dose range of 500 to 2000 mg/day for the treatment of various neuropsychiatric disorders. Hence this dose range was used in this study to estimate the plasma and CSF concentration achievable. VPA has highly variable volume of distribution (Vd, 0.1 – 0.4 L/kg) which contributes to a high degree of variation in the plasma and CSF concentration of VPA. Considering the dose range of VPA used and the variable Vd, this study estimated the plasma concentration of VPA to range from 170 to 7000 µM (figure 1). The CSF concentration of VPA is generally accepted to be ∼ 10% of plasma concentration based on which this study estimated the CSF concentration of VPA to range from 17 to 700 µM (figure 1). At the higher Vd (>10 L), the plasma concentration achieved was within the safety limits (170 to 700 µM in plasma and 17 to 70 µM in CSF), however at lower Vd (<10L), the plasma or CSF concentration achieved was of concern especially while using VPA dose of over 1000 mg/day. While plasma concentration achieved at very low Vd (< 2L) was of concern even at the VPA dose of 500 mg/day (figure 1). Considering these observations, it is trivial to assess the Vd of VPA in each patient to ensure optimal benefits from the treatment and personalise the dose titration. Also in patients with very low Vd the dose of VPA should not exceed beyond 500 mg/day. To further assess the relevance of this highly variable drug concentration achieved in the patients to its therapeutic safety, this study identified all potential targets of VPA and estimated its specific affinity.

**Figure 1:**
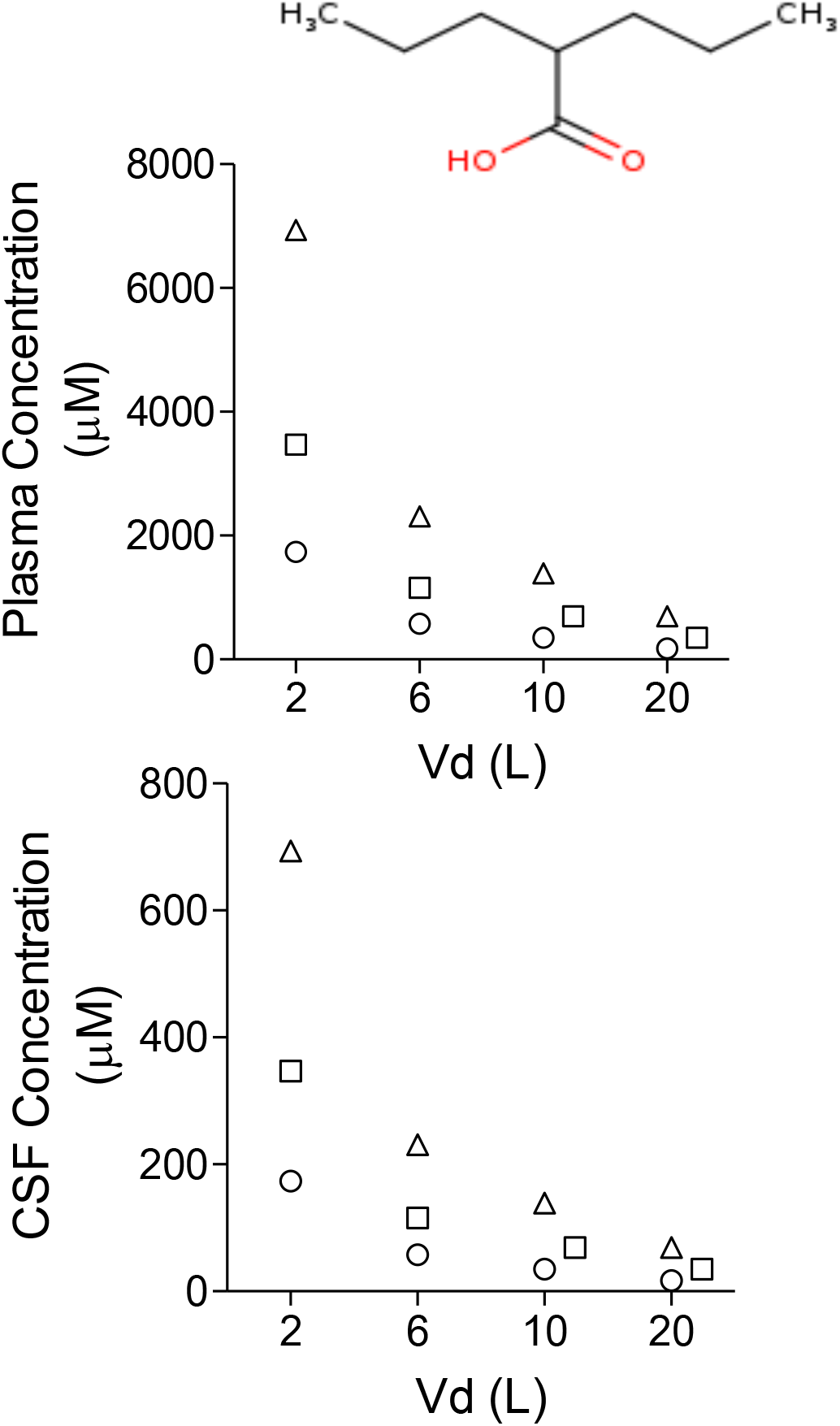
Plasma and CSF concentration of valproic acid (VPA) achieved at variable volume of distribution (Vd, Liters (L)). Inset is the structure of VPA.

The affinity of VPA with its predicted targets, ranged from 2.9 to 52.1 mM (figure 2). The highest affinity (< 5 mM) of VPA was observed against the following targets: Leukocyte common antigen (PTPRC), Lysine-specific demethylase 5C (KDMSC), GABA-B receptor (GABBR1), Histone deactylase 1 (HDAC1), Betaine transporter (SLC6A12), Voltage-gated calcium channel alpha2/delta subunit 1 (CACNA2D1), Cytochrome P450 19A1 (CYP19A1), Niemann-Pick C1-like protein 1 (NPC1L1), Acyl coenzyme A:cholesterol acyltransferase (CES1), Metabotropic glutamate receptor 5 (GRM5), and Solute carrier family 22 member 12 (SLC22A12) (figure 2). The major expression pattern of these high affinity targets in humans is summarised in table 1. The affinity of two major metabolites of VPA (i.e., 2-en-VPA and 3-keto-VPA) were also assessed against these high affinity targets and are summarised in figure 2. The affinity of 3-keto-VPA against most of these targets was superior to that of VPA and was especially observed to be higher against GABBR1, SLC6A12, CYP19A1 and GRM5 (figure 2). While the affinity of 2-en-VPA was inferior to VPA against most targets except GRM5 and SLC22A12 (figure 2).

**Figure 2:**
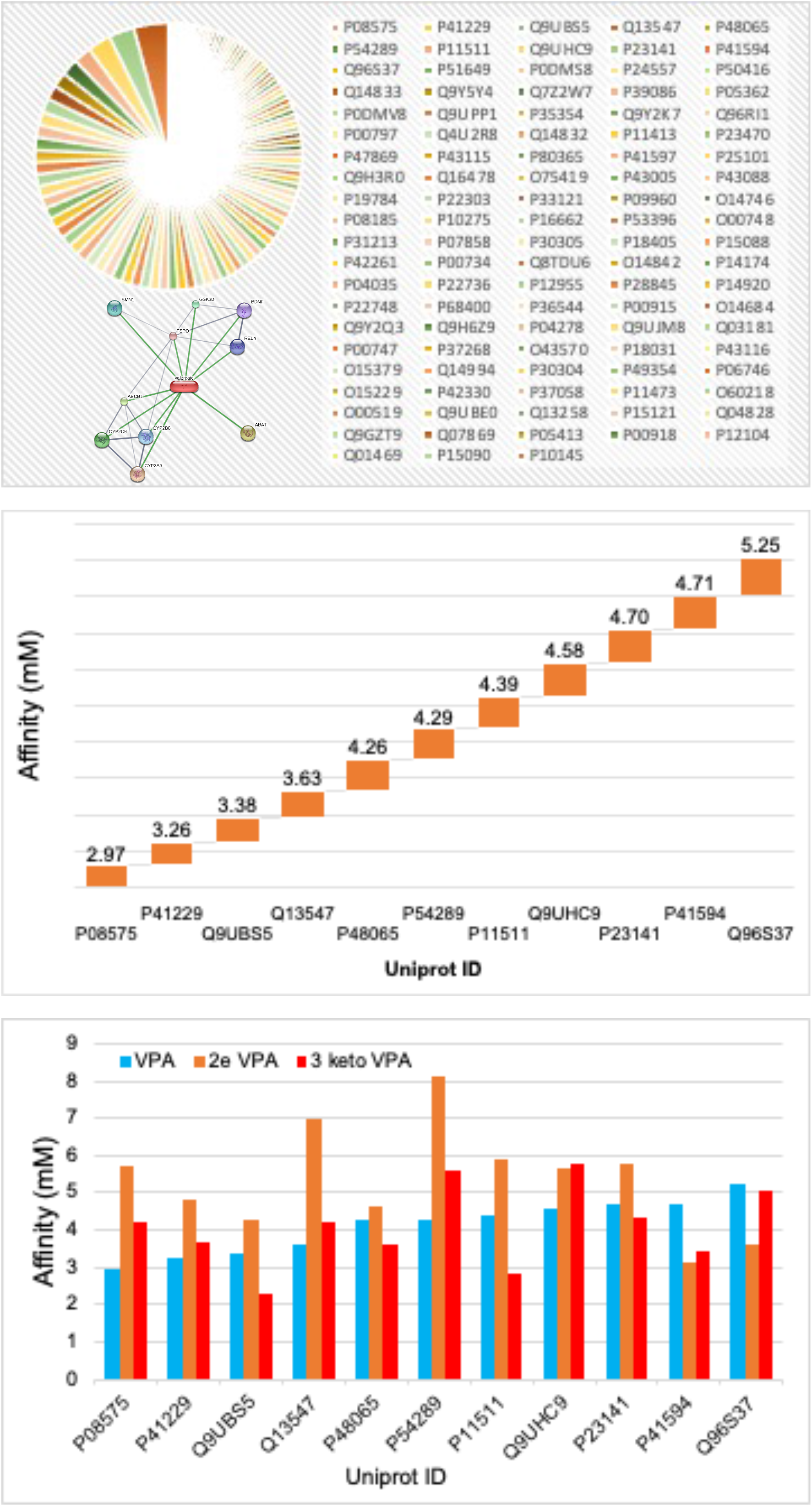
Affinity of valproic acid (VPA) against its human specific targets. Top panel shows affinity of VPA (2.9 to 52.1 mM) to all human specific targets (represented by their uniport ID) extracted from the SwissTargetPrediction server and inset image shows VPA targets identified in the STITCH database. The middle panel shows the high affinity (affinity < 5mM) targets of VPA. The bottom panel shows the comparative affinity of the VPA and its major metabolites (2-en-VPA and 3-keto-VPA) against the high affinity targets.

**Table 1:** Protein expression pattern of high affinity targets of VPA in humans.

| Target | Short name | Uniprot ID | Major Expression |
| --- | --- | --- | --- |
| Leukocyte common antigen | PTPRC | P08575 | Skin, Appendix, Spleen, Lymph node, Tonsil, Bone marrow |
| Lysine-specific demethylase 5C | KDM5C | P41229 | Skeletal muscle, Thymus, Ovary, Bone marrow |
| GABA-B receptor (by homology) | GABBR1 | Q9UBS5 | Cerebral cortex, Cerebellum, Hippocampus |
| Histone deactylase 1 | HDAC1 | Q13547 | Thyroid gland, Bronchus, Oral mucosa, Gastro intestine system, Gallbladder, Urinary bladder, Testis, Epididymis, Fallopian tube, Cervix, Placenta, Appendix, Tonsil, Bone marrow |
| Betaine transporter | SLC6A12 | P48065 | Liver, Kidney, Cerebral cortex |
| Voltage-gated calcium channel alpha2/delta subunit 1 | CACNA2D1 | P54289 | Skeletal muscle, Hippocampus, Ovary, Heart muscle, Soft tissue, Skin |
| Cytochrome P450 19A1 | CYP19A1 | P11511 | Placenta |
| Niemann-Pick C1-like protein 1 | NPC1L1 | Q9UHC9 | Duodenum, Small intestine, Liver, Appendix |
| Acyl coenzyme A:cholesterol acyltransferase | CES1 | P23141 | Nasopharynx, Bronchus, Lung, Liver, Gallbladder |
| Metabotropic glutamate receptor 5 | GRM5 | P41594 | Cerebral cortex, Hippocampal formation, Basal ganglia, Amygdala |
| Solute carrier family 22 member 12 | SLC22A12 | Q96S37 | Kidney |

The affinity of VPA against its selective targets and its plasma or CSF concentration values was used to compute the concentration affinity (CA) ratio, as CA ratio is helpful in analysing the therapeutic safety of a drug. The CA ratio of VPA against each of its high affinity target is shown in figure 3. The CA ratio of VPA against all of its high affinity target was observed to be greater than 0.8, indicating potentially significant modulation of these targets by VPA and hence all these high affinity targets are relevant to the pharmacodynamic effects of VPA. Of specific interest among these targets are PTPRC, KDM5C, GABBR1 and HDAC1 due to their CA ratio being higher than 1 (plasma) or 0.1 (CSF) (figure 3). The general observation being VPA dose of < 1000 mg/day is safer when the Vd is > 10 L while the VPA dose of 2000 mg/day should only be considered in patients when Vd is > 15 L. The VPA dose in patients with Vd < 2L should be restricted to </= 500 mg/day.

**Figure 3:**
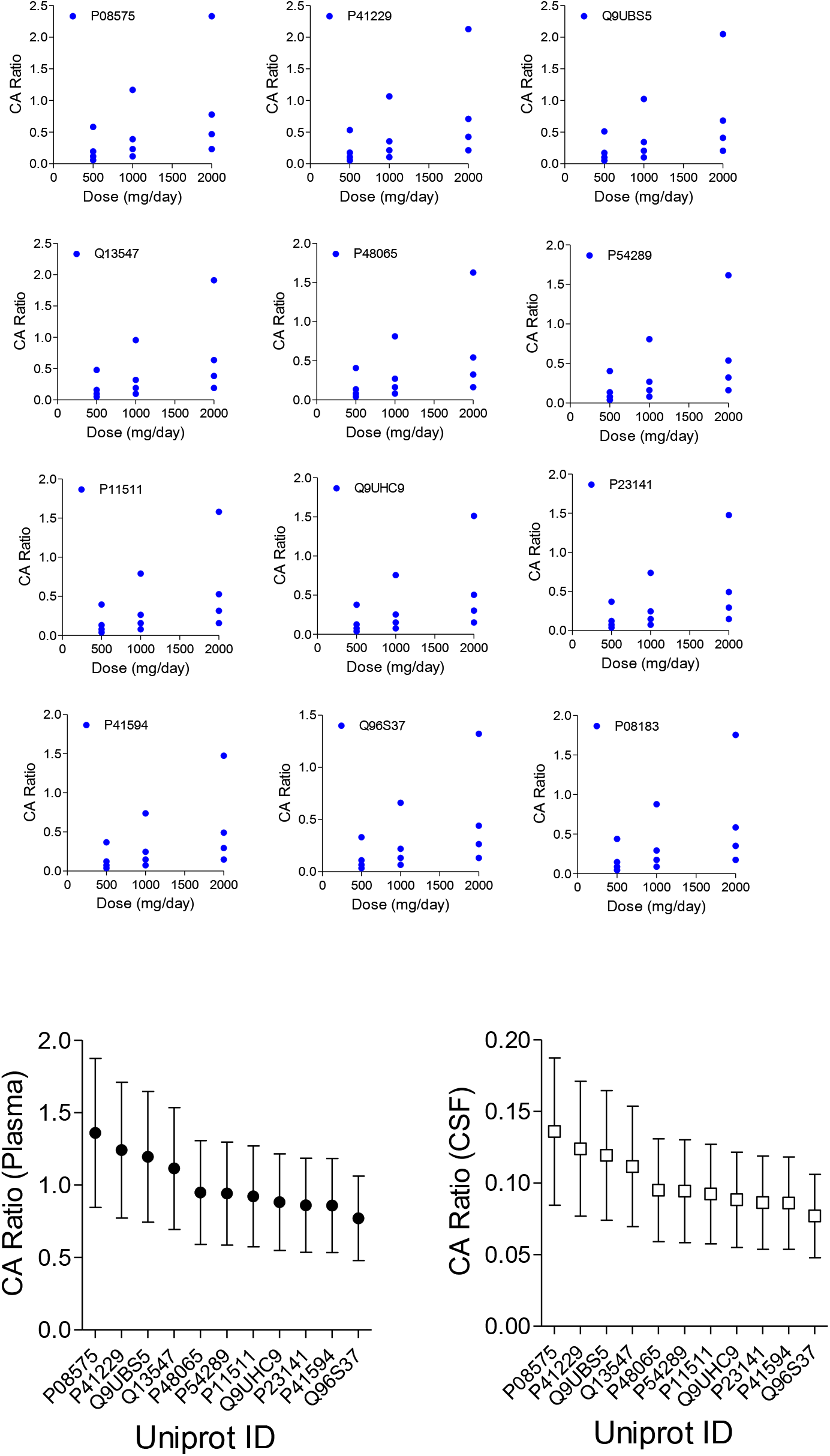
The concentration affinity (CA) ratio of VPA (500 to 2000 mg/day) with all its high affinity targets at variable volume of distribution (2 to 20 litres). The bottom graphs show the average (mean ± SD, n=12) plasma and CSF CA ratio of VPA against high affinity targets (represented by their Uniprot ID).

## Discussion

To understand the pharmacological basis of VPA therapeutic effects, this study assessed the concentration of VPA achievable and correlated it with affinity of VPA against all its potential human specific targets. The plasma (170 to 7000 µM) and CSF (17 to 700 µM) concentrations estimated in this study are consistent with several previously published clinical studies.^[3-5, 16]^ The therapeutic range of VPA in plasma is reported to be 170 to 800 µM and concentration of VPA above 1050 µM is reported to generate undesired effects and potentially saturates protein binding.^[3-5, 16, 17]^ Considering the therapeutic range of VPA together with the plasma or CSF concentration of VPA achieved with the clinically recommended dose of 500 to 2000 mg/day, it is evident that the probability of achieving supra therapeutic range of VPA is very likely. About 2/3^rd^ of plasma or CSF concentration of VPA observed in this study was above the therapeutic range of VPA. An important factor contributing to highly variable plasm or CSF concentration is the wide range of volume of distribution (Vd, 0.1 to 0.4 L/kg) observed in humans.^[3, 5]^ Besides VPA is well absorbed from the GI tract and also easily crosses the blood brain barrier.^[3, 16, 18]^ Hence in the opinion of this study personalised dose titration of VPA will be necessary based on the Vd value of VPA for each patient. Considering the observations from this study, it is suggested that patients (∼ 60kg body weight) can be categorised into three groups based on Vd values i.e., low Vd (<2L), medium Vd (6-10 L) and high Vd (>10L). The VPA recommended dose should be aligned with the Vd values as follows: low Vd patients (</=500mg/day), medium Vd patients (500 – 1000 mg/day) and high Vd patients (<2000mg/day).

This study also assessed all potential targets of VPA. The clinical relevance of the VPA targets was assessed on the basis of its affinity and concentration affinity (CA) ratio^[9]^ against each of the specific targets. All together 110 targets of VPA were identified in this study with affinity values ranging from 2.97 to 52.10 mM. The subsequent analysis in this study was restricted to targets with affinity values < 5mM as the pharmacological interactions with targets having affinity values > 5mM would be likely only at supra therapeutic range of VPA and this is paraphs a major limitation of this study. However two of the targets (CES1 and SLC22A12) with affinity of 4.7 and 5.2 mM are likely to contribute to undesired effects (described latter) of VPA^[2, 7]^ and hence the rationale for focusing on targets with affinity values < 5mM for VPA dose optimisation. Of the 11 targets with affinity < 5mM, only four (GABBR1, SLC6A12, CACNA2D1 and GRM5) were observed to be highly expressed in CNS and hence directly relevant to the therapeutic effects of VPA. The therapeutic efficacy of VPA is attributed to the inhibition of GABA transaminase and succinate semialdehyde dehydrogenase and hence enhancing GABA levels in CNS.^[3-5]^ However in this study the affinity of VPA against GABA transaminase (affinity = 6.1 mM) and succinate semialdehyde dehydrogenase (affinity = 5.7 mM) was higher than 5 mM. Hence in the opinion of this study the anticonvulsant efficacy of VPA is primarily mediated through its effect on GABBR1 (affinity = 3.3 mM), SLC6A12 (affinity = 4.2 mM), CACNA2D1 (affinity = 4.2 mM) and GRM5 (affinity = 4.7 mM) targets. Interestingly the two major metabolites of VPA were also observed to effect these high affinity targets, and hence are likely to influence the overall anticonvulsant pharmacology of VPA. This novel insight into the anticonvulsant property of VPA observed in this study is not previously reported. Besides this several factors modulated through epigenetic mechanisms are also reported to be involved in therapeutic efficacy of VPA.^[3, 4, 6]^ Consistent with several such reports, in this study VPA was observed to have high affinity against histone deacetylase 1 (HDAC1, affinity = 3.6 mM) and Lysine-specific demethylase 5C (KDM5C, affinity = 3.2 mM). Both HDAC1 and KDM5C are involved in modulation of histones and influence epigenetic physiology at several organ systems including CNS.^[3, 4, 6]^ Hence HDAC1 and KDM5C can indirectly influence the anticonvulsant pharmacology of VPA. The CA ratio of VPA against its high affinity targets also revealed further insight into the drug dose at which an optimal therapeutic effects can be achieved. While the CA ratio of all high affinity targets ranged from 0.3 to 2.33 (for plasma) the following four targets showed CA ratio of over 1: PTPRC, KDM5C, GABBR1 and HDAC1, indicating their preferential targeting by VPA. All these four targets seem to be directly or indirectly linked to the VPA therapeutic effects. PTPRC is expressed on microglia cells and has an essential immune regulatory role, the direct relevance of which in anticonvulsant pharmacology is continued to be explored.^[19, 20]^ While the modulation of GABBR1 suggest the direct role of VPA in anticonvulsant pharmacology which is paraphs achieved in synergy with effects of VPA on SLC6A12, CACNA2D1 and GRM5. The anticonvulsant pharmacology of VPA is further supplemented by epigenetic effects of VPA mediated through KDM5C and HDAC1. Some of the high affinity targets of VPA are likely to contributes to its undesired pharmacological effects i.e., CNS oedema and hepatotoxicity.^[2, 7, 16]^ The oedema development is attributed to hyperammonaemia consequence to hepatic failure. High ammonia levels interfere with the TCA cycle and promote neuronal death. It is likely that inhibition of SLC22A12 by VPA, which is essential for renal reabsorption of urates is responsible for hyperammonaemia. While the inhibition of CES1 (liver enzyme responsible for detoxification process) by VPA may further aggravate hyperammonaemia by triggering hepatotoxicity. Again these novel target insight into the undesired pharmacology of VPA is not previously reported. In summary this study offers a novel insight into the anticonvulsant and undesired pharmacology of VPA by specifically identifying the targets involved and recommends an evidence based approach to personalise dose titration of VPA to achieve optimal therapeutic benefits.

## Acknowledgement

Research support from University College Dublin-Seed funding/Output Based Research Support Scheme (R19862, 2019), Royal Society-UK (IES\R2\181067, 2018) and Stemcology (STGY2917, 2022) is acknowledged.

